# Environmental Heterogeneity Altered the Growth Fitness of Antibiotic-Resistant Mutants and the Resistance Prevalence in *Escherichia coli* populations

**DOI:** 10.1101/2025.08.18.670914

**Authors:** Chujing Zheng, Yue Xing, Xiaoxi Kang, Yujie Men

**Author notes:** These two authors contributed equally to this work (Chujing Zheng, Yue Xing).

## Abstract

Antibiotic resistance is one of the most critical issues in public health. As antibiotic-resistant bacteria emerge under certain selection pressure, their further proliferation can facilitate the prevalence and maintenance of the antibiotic resistance. Here, we investigated which environmental factors could affect the growth fitness of antibiotic-resistant *Escherichia coli* strains using growth competition assays. We found that when antibiotic resistance selection pressure was removed, lower temperature and nutrient limitations, especially iron (Fe) deficiency, fostered a better fitness to resistant mutants when co-cultivated with the wild type, whereas changes in pH or salinity (Na or K) did not. Moreover, the growth advantage of resistant mutants under the selection pressure was reversed by suboptimal conditions like acidic/basic pH, high K salinity, and Fe limitation. These identified key environmental factors influencing the growth fitness of antibiotic-resistant mutants provide important insights into the prediction and mitigation of antibiotic resistance in heterogeneous environments.

## 1. Introduction

The spread of antibiotic resistance has threatened the effectiveness of antibiotic therapy, drawing increasing attention in scientific and public communities (He et al., 2023; Murray et al., 2022; Peng et al., 2023). Antibiotic resistance could be developed and propagated via vertical (*de novo* mutation) and horizontal gene transfer (HGT). Studies have shown that not only antibiotics but also non-antibiotic chemicals (e.g., pesticides, pharmaceuticals, and heavy metals) introduced to environments via water and biosolids reuse during agricultural practices could facilitate the emergence of antibiotic-resistant bacteria (ARB) or the dissemination of antibiotic resistance via HGT (Phan et al., 2024; Tang et al., 2015; Wang et al., 2020; Wang et al., 2019; Xing et al., 2021; Zhang et al., 2023). This would increase the exposure of ARB to humans via the food chain, posing a potential risk to public health (Becerra-Castro et al., 2015; Bhattacharjee et al., 2024; Phan et al., 2024; Sun et al., 2021).

One important step for ARB to thrive and spread upon emerging is to survive and proliferate in a given environment, where ARB and susceptible strains may co-exist and compete for limited living resources. The proportion of ARB in populations could determine the level of antibiotic resistance in microbial communities (Letten et al., 2021). Mostly, there are fitness costs by possessing the resistance phenotype (Andersson and Hughes, 2010; Hinz et al., 2024; Melnyk et al., 2015). As the environmental conditions favoring the resistance phenotype change or when evolved resistant mutants enter a new environment, indigenous susceptible strains may suppress the proliferation of resistant strains by competing for resources and ecological niches under specific geochemical conditions (Hinz et al., 2024; Kinsler et al., 2020; Klümper et al., 2019). This competition process plays a key role in determining the successful propagation and consequently modulating the prevalence of ARB (Davies et al., 2019; Letten et al., 2021). Therefore, understanding how the competition between ARB and susceptible strains respond to varying geochemical properties is critical for predicting the distribution and prevalence of antibiotic resistance and developing mitigation strategies in heterogeneous environments (e.g., soils).

Recent resistomes studies revealed correlations between environmental factors (e.g., pH and salinity) and the abundance and distribution of antibiotic resistance genes (ARGs) in soil microbial communities (Liu et al., 2023; Tan et al., 2019; Xu et al., 2023), but with some inconsistent findings. Xu et al. (Xu et al., 2023) found that high salinity (as Na) enriched ARGs and mobile genetic elements in soil microbiota, whereas Tan et al. (Tan et al., 2019) found the opposite. This could be due to the difference in the soil composition and soil microbial communities used in those studies. Moreover, variations in the relative abundance of ARGs would not necessarily point to the same change in resistance phenotype. Pure culture studies showed that nutrient limitations induced antibiotic resistance, suggesting increased fitness of resistant mutants under those conditions (Eng et al., 1991). This was supported by one competition study, where the resistant strain gained a higher growth fitness over the wild type under limited carbon sources (Lin et al., 2018). This could be due to changes in metabolic activities and adaptive mutations, but it was unclear whether the adaptive mutations caused the metabolic changes, as the two co-occurred. Therefore, to disentangle the role of individual environmental factors in shaping the growth fitness and modulating the prevalence of resistance in populations, studies under simple and well-controlled conditions are needed.

In this study, we aimed to identify environmental factors that can alter the growth fitness of ARB, hence potentially modulating the spread of antibiotic resistance via the evolve-proliferate (i.e., genotype-phenotype-fitness) route. We conducted growth competition assays between resistant and susceptible strains in co-cultures. We comprehensively investigated common geochemical parameters, including pH, temperature, salinity (as Na and K), and key nutrients, at relevant levels found in heterogeneous environments such as soils. Mimicking the evolve-proliferate route, we followed the continuity of previous evolution studies and used resistant *Escherichia coli* mutants evolved under a long-term exposure to an environmentally relevant selection pressure, i.e., sub-lethal streptomycin and a mixture of pesticides (Xing et al., 2021). The genotypes, i.e., single-nucleotide polymorphism (SNP) and insertions/deletions, of investigated resistant *E. coli* mutants have been identified (Xing et al., 2021). Competition assays were conducted between resistant mutants and the streptomycin-sensitive wild type in co-cultures under growth conditions with only one varying parameter. The fitness of resistant mutants relative to the wild type was determined using relative abundances measured by SNP-based genotyping assays. Environmental factors altering the fitness of resistant mutants were then identified by comparing the change in fitness under different conditions.

## 2. Results and Discussion

### 2.1 Nutrient deficiency and low temperature weakened the growth disadvantage when co-cultivated with the wild type without the antibiotic-resistance selection pressure

Most antibiotic resistance mutants are associated with a fitness cost, which can be offset by the resistance phenotype under the selection pressure (e.g., presence of antibiotics). Thus, the removal of resistance selection pressure could reverse the growth advantage and dominance of resistant mutants over the wild type (Andersson and Hughes, 2010). However, given the heterogeneity of certain environments and the change of residing environments after the mutant emerged, one key question is whether there are specific environmental conditions that can reverse the growth disadvantage of resistant mutants in the absence of selection pressure. Here, we tackled this question using competition assays in co-cultures of streptomycin-resistant and susceptible *E. coli* strains under various growth conditions. We specifically looked at the role of environmental factors by comparing the relative fitness of resistant mutants after a short period (30 generations) (Xing et al., 2021), so that the effect of compensatory mutations on fitness cost could be minimized (Andersson and Hughes, 2010).

Results showed that all tested environmental conditions did not alter the dominance of the susceptible wild type over the resistant mutant in the absence of selection pressure, i.e., with the relative fitness (F) of the resistant mutant below 1 (**Fig. 1**). Despite no effect of pH and salinity (Na or K), lowering the temperature slightly restored the growth advantage of four resistant strains (S1, S2, M1, and M2), elevating their relative abundances by one order of magnitude: from ∼ 1% (F = 0.85) under the optimal temperature at 37°C to ∼ 11% (F = 0.93) at 20°C (**Fig. 1**). Cold stress was reported to repress the expression of ARG and virulence factors (Grosso-Becerra et al., 2014; Zarzecka et al., 2022). Similarly, the gene mutations conferring streptomycin resistance (Table 1) could be down-regulated under low temperature, thus reducing the fitness cost of resistant mutants.

**Figure 1.**
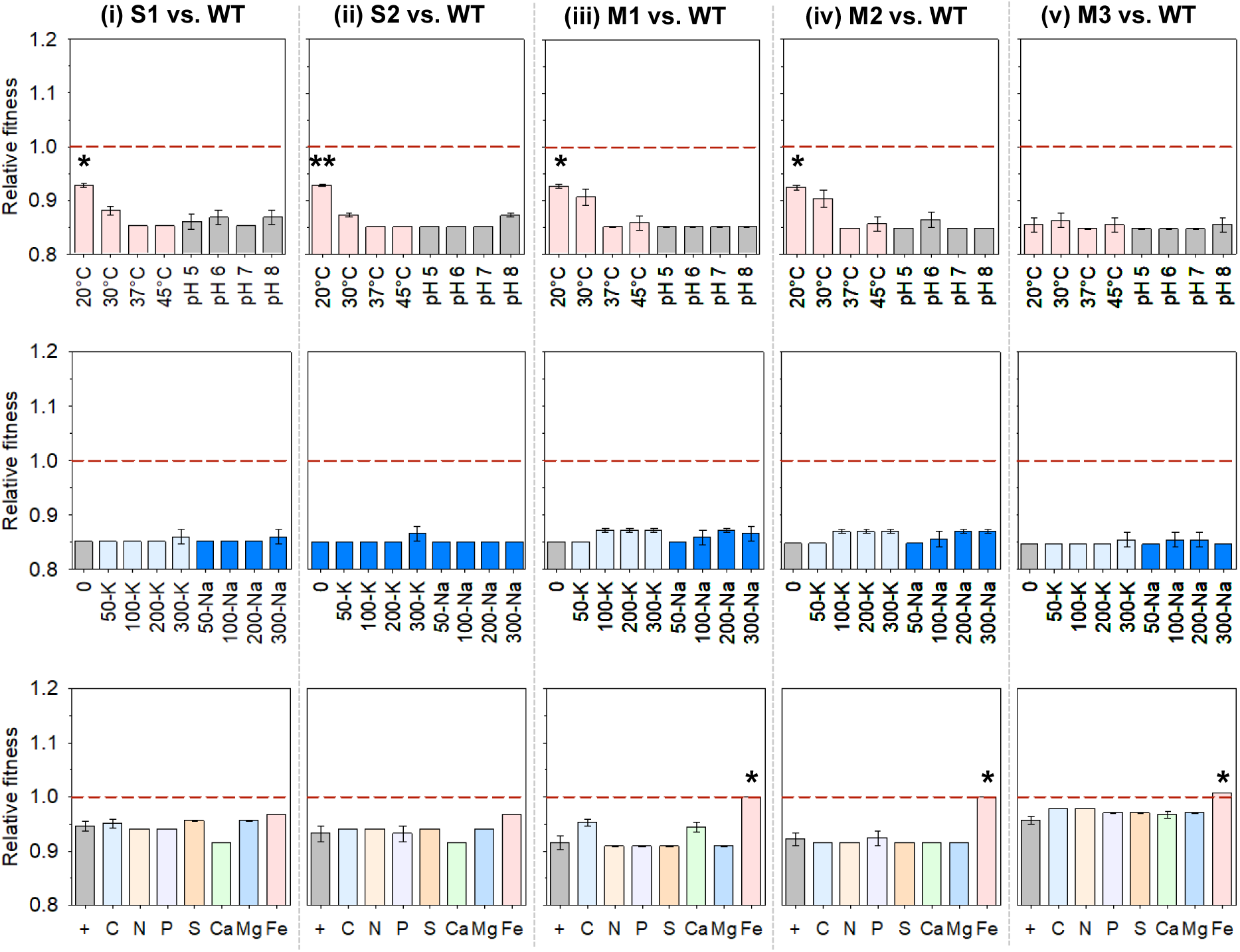
The relative fitness of each resistant mutant, (i) S1 (ii) S2 (iii) M1 (iv) M2 (v) M3, versus the wild type (WT) under different growth conditions **in stress-free media** (top: temperature and pH in LB medium; middle: additional salinity as NaCl or KCl in mM in LB medium at pH 7, 37 °C; bottom: elemental nutrient limitations in M9 medium, “+”: the original M9 minimal medium with 0.4% glucose, at pH 7, 37 °C; Welch’s t-test for differences between suboptimal conditions and the respective optimal condition of the tested parameter, *: *p* < 0.05, **: *p* < 0.01, n = 3).

**Table 1.** *E. coli* strains used in this study.

| Strains | Resistance genotype | Fold change of MIC* |
| --- | --- | --- |
| Wild type | *E. coli* strain K-12 (ATCC 10798, streptomycin-sensitive) | 1 |
| S1 | *rpsL* (Arg86Ser), *rsmG* (Trp150fs), *dsbC* (Val172Glu) | 40 |
| S2 | *rpsL* (Arg86Ser), *rsmG* (Trp150fs) | 25 |
| M1 | *glnE* (Ala423Val), *yaiW* (Phe183Ile, Gln186Asp, His187fs) | 4 |
| M2 | *nuoG* (Ser548**) | 3 |
| M3 | *glnE* (Ala423Val), *nuoG* (Glu10**), *sbmA* (Glu282**) | 3 |

\*The ratio of MIC to MIC<sub>0</sub>, where MIC<sub>0</sub> is the MIC of streptomycin for the wild type, 8 mg/L.
\*\*Stop codon

Although the wild type remained dominant under all investigated nutrient conditions, the switch from the nutrient-rich condition to oligotrophic conditions (i.e., M9 minimal medium with varying elemental nutrient levels) diminished the fitness effect for resistant strains in general, with an increase of relative fitness from 0.85 to above 0.9 but still below 1 (**Fig. 1**). This is similar to what has been observed in streptomycin-resistant *Salmonella enterica* var. Typhimurium LT2 mutants, which exhibited relatively faster growth rates when grown on poorer carbon sources (Paulander et al., 2009). Moreover, the Fe limitation condition further reversed the growth disadvantage of the mildly resistant mutants (M1 – 3) and made them as fit as the wild type with the relative abundance stayed unchanged (∼ 1.0) from the beginning (**Fig. 1**). A titration experiment showed that lowering the free Fe availability to 34 μg/L doubled the relative abundance of M1. When the free Fe was further lowered to 34 ng/L, the mildly resistant mutant exhibited a similar growth fitness to the wild type (**Fig. S1**). Such an effect was not observed for the two strongly resistant strains, suggesting a critical role played by the mutant genotypes and resistance mechanisms. The mutants with mild streptomycin resistance possessed mutations on off-target genes involved in cell transport and metabolic activities (*glnE*, *yaiW*, *sbmA*, and *nuoG*), whereas the strongly resistant mutants had mutations on the streptomycin-target genes (e.g., *rpsL*). The off-target mutations might provide higher resilience to environmental stresses in addition to antibiotics, while the target mutations offered a strong resistance to antibiotics at a higher cost of growth fitness when co-cultivated with the wild type.

The above findings imply that the elimination of antibiotic resistance selection pressure, including antibiotics and non-antibiotic chemicals known to induce resistance, can be an effective way to suppress the proliferation and prevalence of emerged resistant mutants in varying receiving environments of ARB. Nevertheless, special cautions should be made for low temperature and nutrient-limiting (especially Fe) conditions in some aquatic environments like drinking water sources and drinking water storage tanks, as those conditions tend to prevent resistant mutants from being outcompeted by susceptible strains, promoting the survival of resistant strains and maintaining their relative abundance. It could also explain the resilience of ARB in drinking water systems reported in other studies (Lin et al., 2018; Zhang et al., 2018).

### 2.2 Suboptimal pH and high K salinity reversed the growth advantage of resistant strains under the selection pressure

Besides controlling environmental factors to suppress the proliferation of ARB in receiving environments without the selection pressure, it is also critical to identify key parameters that can alter the growth fitness of resistant mutants as they emerge over the evolutionary course of antibiotic-resistance selection in antibiotic-laden environments. We conducted competition assays under various environmental conditions, but with the same selection pressure as the resistant mutant was isolated. As expected, with the sustained selection pressure, the growth fitness of all resistant mutants relative to the wild type was greater than 1 under optimal growth conditions (37 °C, pH=7, no salts added in the LB medium), indicating the selection direction toward antibiotic resistance. This growth advantage of the resistant mutant (with F = 1.05 – 1.1, ca. 82 – 96% relative abundance of the resistant strain) over the wild type under the selection pressure was successfully maintained at suboptimal temperatures and higher external Na concentrations (**Fig. 2**), suggesting that altering those environmental factors could not interfere with the resistance selection. Nonetheless, unfavorable pH (both acidic and basic) and high external K concentrations (higher than 100 mM) completely reversed the dominance of the resistant mutant (F < 0.9, with a relative abundance < 10%) in the co-culture under the selection pressure (**Fig. 2**). The slightly acidic pH (5 – 6) substantially overturned the selection of all streptomycin-resistant strains with either strong or mild resistance, while the effect of the slightly basic pH (8) was not as strong as the acidic pH for the strongly resistant strains compared to the mildly resistant ones (**Fig. 2**). The dominance of the strongly resistant mutant (S1) was also overturned when co-cultivated with the mildly resistant mutants under suboptimal pH and high external K concentrations (**Fig. S2**), suggesting a general decrease in the overall streptomycin resistance of populations in either the early (consisting of both resistant and sensitive strains) or late (only consisting of mutants with different resistance levels) stages of resistance evolution (Xing et al., 2021).

**Figure 2.**
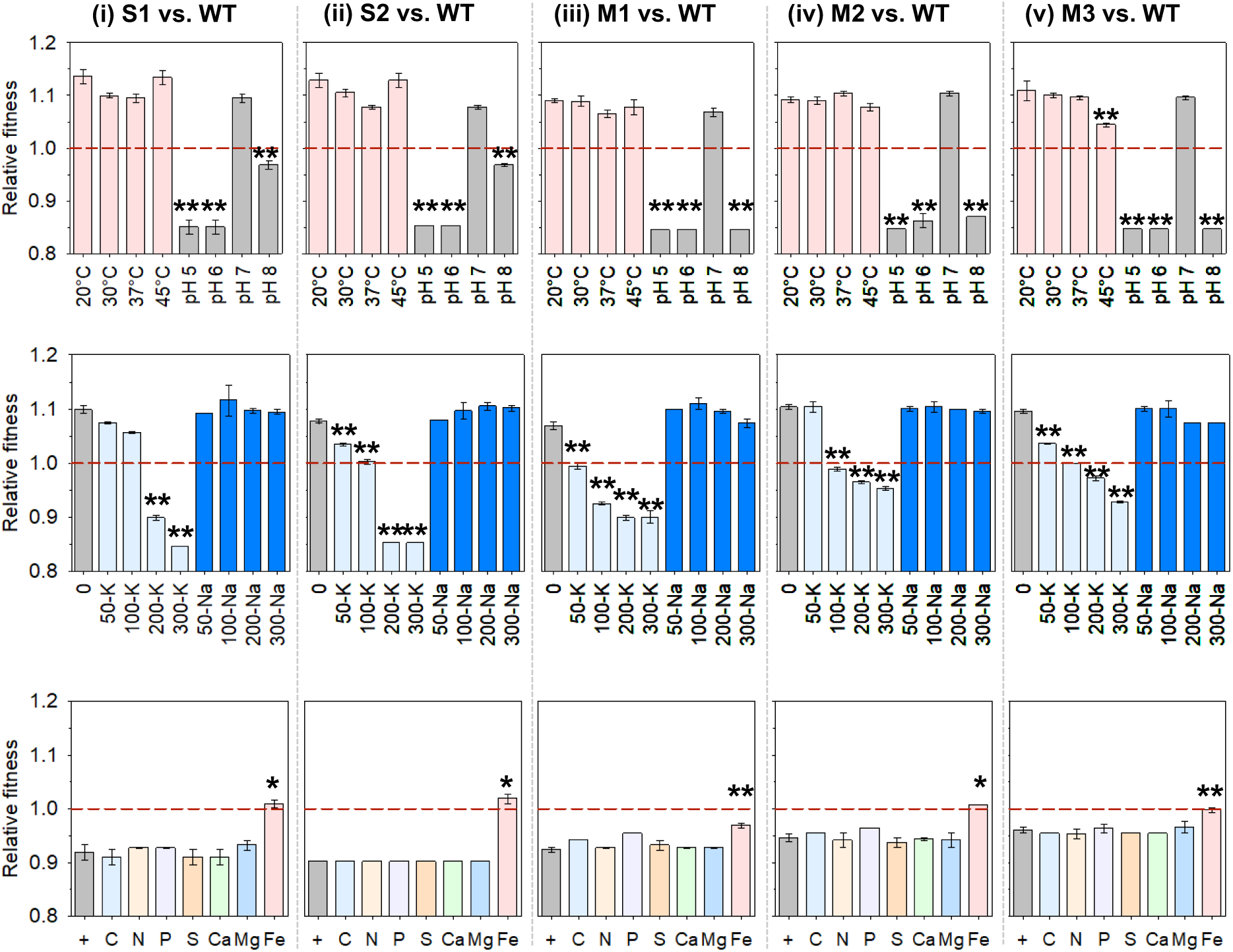
The relative fitness of each resistant mutant, (i) S1 (ii) S2 (iii) M1 (iv) M2 (v) M3, versus the wild type (WT) under different growth conditions **with the selection pressure** (top: temperature and pH in LB medium; middle: additional salinity as NaCl or KCl in mM in LB medium at pH 7, 37 °C; bottom: elemental nutrient limitations in M9 medium, “+”: the original M9 minimal medium with 0.4% glucose, at pH 7, 37 °C; Welch’s t-test for differences between suboptimal conditions and the respective optimal condition of the tested parameter, *: *p* < 0.05, **: *p* < 0.01, n = 3).

The uptake and efficacy of streptomycin is pH dependent. The increase of pH can enhance the electrical component of the proton motive force, hence the uptake of aminoglycoside antibiotics including streptomycin, and vice versa (Donovick et al., 1948; Webster and Shepherd, 2022). Thus, one possible explanation for what was observed under acidic pH is that the low pH could inhibit the uptake of streptomycin for all resistant and sensitive strains. This diminished the effect of streptomycin, making it similar to the stress-free medium-only condition (**Fig. 1**), where the wild type had a higher growth fitness than the resistant mutants. It is also supported by the growth curve of the resistant mutants and the wild type under different pH levels with and without the selection pressure. Under pH 5, the resistant strains (S1 and M1) showed slower growth and lower maximum cell density than the wild type, the same as those without the selection pressure at pH 5 (**Fig. 3A & B**), but they exhibited better growth with the selection pressure than without at pH 7 (**Fig. 3E & F**). Thus, this altered relative growth fitness of resistant and susceptible strains could be due to the decreased streptomycin potency at low pH. On the other hand, at the basic pH, streptomycin uptake would be enhanced and make the wild type more sensitive in the presence of the selective pressure, as it lacks a resistance mechanism. However, the relative growth fitness of the wild type increased at pH 8, suggesting that higher pH had a more negative growth effect on the resistant strains, especially the mildly resistant ones, hence decreasing their relative fitness.

**Figure 3.**
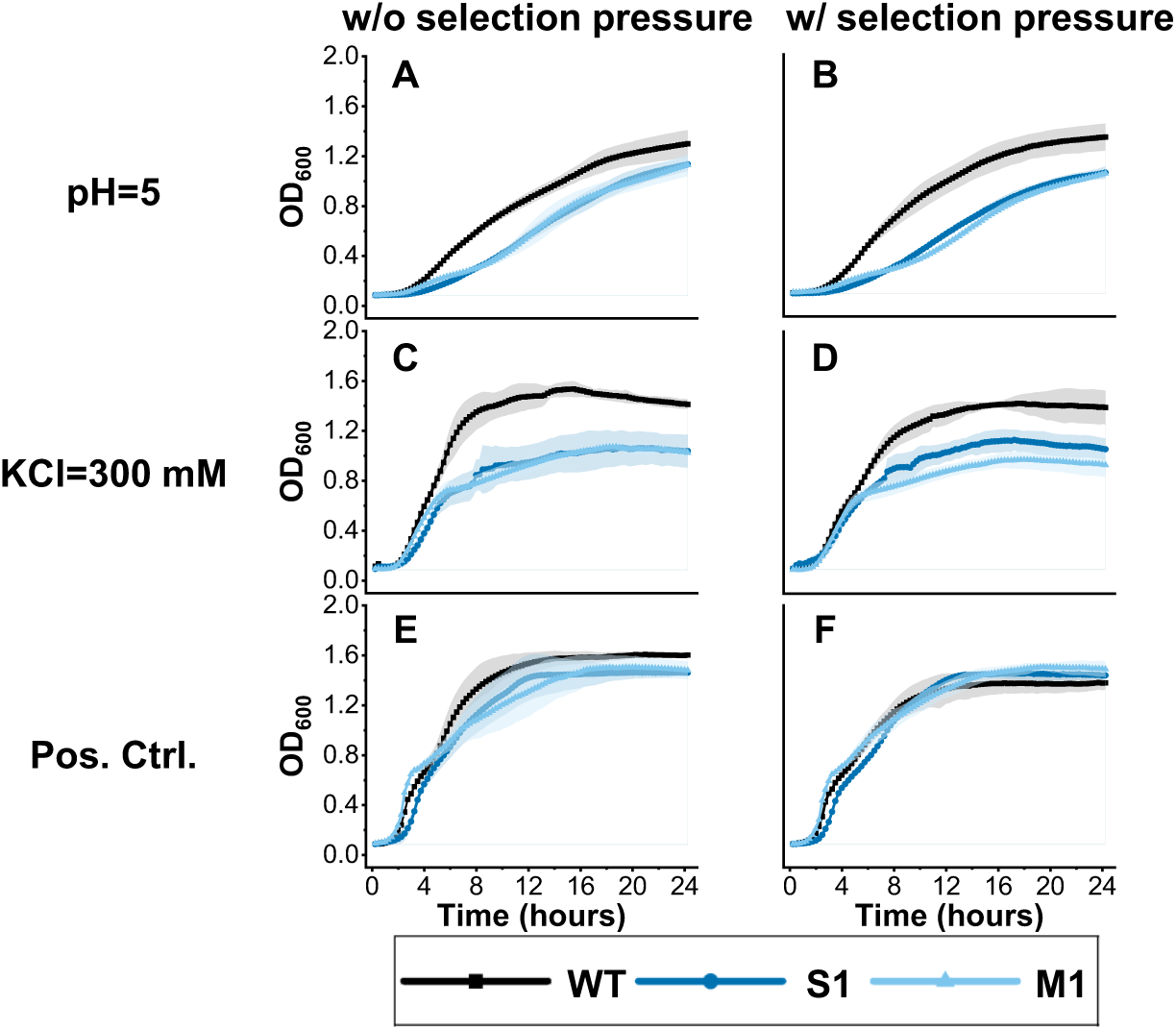
Growth curves of the wild type, S1, and M1 under low pH (A & B) and high external K (C & D) in the absence (left panels) and presence (right panels) of the selection pressure, compared to the positive control in standard LB medium at pH 7 (E & F) (Curves represent the average OD_600_ of 4 – 6 replicates and shades represent the standard variation).

The reversed selection for streptomycin-resistant strains under high K concentrations was also likely due to a decrease in streptomycin potency, similar to the low pH condition. One study reported that streptomycin potency was dependent on a K^+^ efflux channel (MscL), which gates to release solutes, including K^+^, in response to a hypo-osmotic down shock (Iscla et al., 2014). According to that study, streptomycin could induce MscL, and the open MscL pore could serve as a pathway for streptomycin into the cell cytoplasm (Iscla et al., 2014). High external K^+^ concentrations likely suppressed the expression of MscL, thus reducing streptomycin potency to the wild type under the selection pressure and amplifying the fitness cost of resistant mutants. Alternatively, it may be explained by the increase of membrane potential (depolarization) caused by high external K^+^. The high external K^+^ can induce another mechanosensitive channel MscK (Li et al., 2002). The opening of MscK channel led to membrane depolarization and an increase in membrane potential (Stautz et al., 2021), the same as the effect of low pH (Felle et al., 1980), which hindered the uptake of streptomycin (Donovick et al., 1948). Therefore, the high external K^+^ undermined the potency of streptomycin, reversing the relative growth fitness of the resistant mutants and the wild type in the presence of the selection pressure. This is also reflected by the growth curves of resistant mutants with the addition of 300 mM KCl with and without the selection pressure (**Fig. 3C & D**), which were similar to those at pH 5 (**Fig. 3A & B**) in comparison to the positive control (**Fig. 3E & F**), suggesting the same consequence from hindered streptomycin update under those two conditions.

### 2.3 Fe deficiency favored the selection of streptomycin-resistant strains, while the limitation of other nutrients reversed the selection direction under the selection pressure

The resistant mutants originally evolved in a nutrient-rich medium (LB). Switching the growth condition from nutrient-rich to nutrient-poor/limited conditions, except for Fe-limitation, overturned the resistance selection direction, even under the selection pressure (**Fig. 2**). The relative growth fitness of all resistant strains in the original M9 medium (0.4% glucose) decreased below 1.0 (0.9 – 0.95) compared to 1.1 and above in the original LB medium under the optimal pH and temperature. It implies that the increased fitness cost under nutrient-poor conditions outweighed the growth benefit provided by the resistance mutation in the resistant strains, making them outcompeted by the wild type. Further lowering the concentration of individual elemental nutrients (C, N, P, S, Ca, and Mg) did not contribute more. In contrast, Fe deficiency significantly restored the growth advantage of resistant strains over the wild type (**Fig. 2**). This is the same as in the medium without the selection pressure.

Studies have shown that Fe availability may affect the efficacy of antibiotics, and how it affects the antibiotic potency could be antibiotic-specific and via complex mechanisms (Ezraty and Barras, 2016). Since Fe is involved in many metabolic processes, a low external Fe availability might induce Fe transport, which potentially facilitates the uptake of certain antibiotics, hence antibiotic potency (Mochizuki et al., 1988). Under Fe deficiency, bacteria may produce siderophores and use them as an efficient Fe carrier to transport Fe into cells. Meanwhile, antibiotics could form conjugates with siderophores and get co-transported into cells as a “Trojan Horse” strategy to increase antibiotic sensitivity (Rayner et al., 2023). A naturally produced siderophore-antibiotic conjugate is the aminoglycoside-danoxamine, salmycin (Rayner et al., 2023). Researchers also observed Fe-mediated abiotic conjugation of siderophores and antibiotics (Caradec et al., 2023). It suggests that this siderophore-antibiotic co-transport mechanism enhanced the efficacy of streptomycin to the wild type strain in the co-culture under Fe limitation conditions. Thus, the sublethal level of streptomycin could have a higher inhibition on the growth of the wild type. In contrast, it did not affect the growth of the resistant strains that can resist a much higher level of streptomycin.

### 2.4 Implications and limitations

The results of this study emphasize the importance of environmental factors on the survival and proliferation of ARB in heterogeneous environments. The identified conditions that provide resistant strains with growth advantages over sensitive ones in a population can be used to predict and prevent the outgrowth and prevalence of ARB. These insights apply across environments, including agricultural soils, surface water bodies receiving discharges and treated wastewater, wastewater and activated sludge communities, as well as gut microbiomes in humans and animals. To contain the relative abundance of resistant strains at low levels in an environmental community, the very first and most effective way would be the removal of antibiotic resistance selection pressure, such as antibiotics and non-antibiotic chemicals known to induce resistance. Compared to sensitive microbes in a natural community, the higher fitness cost associated with most resistant strains can limit their outgrowth in the community when the selection pressure is removed. In other words, one important strategy to prevent antibiotic resistance from spreading is to minimize the occurrence in the environment from potential sources and promote proper use of antibiotics to control exposure to humans and animals. Furthermore, when possible, altering environmental conditions to those not favoring the survival and proliferation of resistant strains may also help mitigate the antibiotic resistance risk, for example, switching to acidic or basic pH and maintaining a relatively higher external K^+^ level without adverse effects on soils or plant growth in agricultural environments. Maintaining a sufficient Fe level in soil and gut microbiomes could also prevent resistant mutants from outcompeting the sensitive strains.

It is worth noting that the identified environmental conditions, which altered the relative growth fitness of streptomycin-resistant strains, may be specific to antibiotics, antibiotic resistance genotypes, and bacterial species (Hinz et al., 2024). For example, instead of reducing streptomycin potency and diminishing the fitness advantage of resistant strains, extremely low pH was found to benefit the survival of multidrug-resistant *E. coli* with an MdtEF-TolC multidrug efflux pump (Schaffner et al., 2021). One should exercise caution when extrapolating the findings in this study, which used streptomycin-resistant *E. coli* with specific genotypes to genotypes conferring resistance to antibiotics from other categories than aminoglycosides. Nonetheless, as aminoglycoside antibiotics share similar chemical structures, they may be taken up by cells via similar mechanisms, which could be affected by the same environmental conditions identified in this study, such as pH, external K^+^, and Fe levels. Thus, our findings will more likely apply to other aminoglycoside antibiotics. Moreover, our results suggest that the significantly altered growth fitness under certain environmental conditions could be mostly attributed to the changes in antibiotic uptake, and hence antibiotic efficacy. Conditions impacting the uptake of antibiotics could then be predicted based on the physicochemical properties and transport mechanisms, and those conditions will likely alter (negatively or positively) the relative growth fitness between the resistant and susceptible strains. Meanwhile, given the diversity of antibiotics, bacterial species, and resistance genotypes, extensive and comprehensive studies are needed in the future to identify the environmental factors that impact the relative abundance of specific antibiotic-resistant bacterial strains and resistance genotypes via modulating the relative fitness cost.

We also acknowledge that due to the nearly identical genomes between the streptomycin-resistant and the wild-type strains in co-cultures (only differing by several genetic mutations), we were unable to compare the transcriptional and translational differences between the two strains and demonstrate the mechanisms of growth fitness changes at the molecular level. The phenotypic demonstration by the growth competition assays warrants deeper mechanistic studies using appropriate tools, including fluorescence-activated cell sorting (FACS) and genetic modification, to verify proposed molecular mechanisms.

## 3. Conclusions

In this study, we demonstrated that the fitness landscape and the prevalence of streptomycin-resistant *E. coli* shifted as geochemical conditions changed in heterogeneous environments, regardless of the selective pressure for antibiotic resistance. Generally, resistant strains showed a lower growth fitness than the wild type in environments without antibiotic resistance selection pressure (e.g., antibiotics). Some specific conditions (i.e., suboptimal pH, high external K^+^, and C-limiting conditions) could reverse the growth advantage of resistant strains under selection pressure, while Fe deficiency increased the fitness of resistant strains regardless of selection pressure. These findings provide a better understanding of the potential proliferation and outgrowth of antibiotic resistance, especially for the aminoglycoside antibiotics, under varying geochemical conditions in heterogeneous environments. It highlights the need to consider the impact of environmental heterogeneity on the survival and prevalence of ARB when predicting and mitigating antibiotic resistance risks in natural matrices. Future studies may focus on the mechanistic understanding using advanced molecular approaches and expanding the relative fitness study to different antibiotic categories, resistance genotypes, and bacterial species, which may be found in soils, plant tissues, and gut microbiota.

## 4. Materials and Methods

### 4.1. Bacterial strains

Bacterial strains used in this study included a wild-type *E. coli* strain K-12 (ATCC 10798) that is sensitive to streptomycin and five streptomycin-resistant mutants with strong or mild resistance levels. The resistant mutants were isolated from populations of the same wild type strain after co-exposure to sublethal streptomycin (1/5 of the minimum inhibitory concentration, MIC, of the wild type) and a mixture of pesticides at environmentally relevant levels (20 – 2000 μg/L total) for ∼ 500 generations. Details of the evolution experiment, where the resistant mutants were obtained, can be found in our previous studies (Xing et al., 2021; Xing et al., 2020). Genetic mutations conferring streptomycin resistance were also identified (Xing et al., 2021). The streptomycin resistance level (in terms of the fold change of MIC for the resistant mutant compared to that for the wild type) and the resistance SNPs of resistant mutants were listed in **Table 1**. Two mutants exhibited more than 25-fold increase in MIC, representing **<u>s</u>**trong resistance (denoted “S1” and “S2”), while the other three had only less than 5-fold increase, representing **<u>m</u>**ild resistance (denoted “M1-3”). The wild type and resistant mutants were archived in aliquoted glycerol stocks at –80 °C. Before the experiment, they were revived from the stock by adding 2 µL of thawed aliquots into 2 mL of fresh Luria-Bertani (LB) media, then incubated overnight at 37 °C at 200 rpm on an orbital shaker. Those revived cultures were referred to as Generation 0 (G0).

### 4.2. Growth conditions

Effects of various environmental factors were investigated using different sets of growth conditions. Each set of growth conditions only included one varying environmental factor, i.e., pH, temperature, Na, K, and other required elemental nutrients (C, N, P, S, Ca, Mg, Fe). All other parameters were set at the default value, i.e., pH = 7, 37°C, no additional salts added, in LB (for pH, temperature, and salinity) or M9 medium (for the limited elemental nutrient conditions). The range of environmental factors was chosen according to those commonly occurring in soil environments, a representative heterogeneous environment, as well as those under which *E. coli* can survive (Kumar and Libchaber, 2013; Nagai et al., 1991; Tan et al., 2020; Zhao et al., 2019). The series of temperatures was realized by incubating the competition co-cultures in different shaker incubators with the temperature set at 20 °C, 30 °C, 37 °C, and 45 °C separately. The pH series (5, 6, 7, and 8) was accomplished by adjusting the pH of LB media with citric acid-sodium citrate buffer solutions for pH = 5 and 6 ± 0.2, Na_2_HPO_4_-NaH_2_PO_4_ buffer solutions for pH = 8 ± 0.2, and no adjustment for pH = 7 ± 0.2. Two representative types of salinity, as Na or K, were tested using a series of concentrations (0, 50, 100, 200, and 300 mM) by adding respective volumes of 4 M KCl or NaCl stock solution into fresh LB media. For the effect of nutrient levels, seven required nutrients, i.e., C, N, P, S, Ca, Mg, and Fe, were tested using the M9 minimal medium with 0.4% (4 g/L) glucose (Sambrook et al., 1989) (see ingredients in **Table S1** of the Supplementary Materials), instead of LB media. The elemental nutrient concentrations in the original M9 minimal salts medium were used as the positive control for comparison. Suboptimal nutrient levels (i.e., nutrient-limited conditions) were chosen (**Table S1**) as the concentration causing 50% decrease (measured as the Optical Density at 600 nm, OD_600_) of the wild type.

Besides the above common geochemical factors, the effect of continuous selection pressure on the proliferation of resistant mutants after they are selected was also investigated. Competition assays were performed with and without the presence of the same selection pressure, which led to the emergence of resistant mutants. Specifically, the selection pressure was of streptomycin at 1/5 of its original MIC for the wild type and a mixture of pesticides at environmentally relevant concentrations. The stock solutions of antibiotics and pesticides were prepared and added to the culture medium in the same way as reported previously (Xing et al., 2021; Xing et al., 2020).

### 4.3. Competition Assays

The growth fitness of resistant mutants under different environmental conditions was examined using competition assays, where two strains were co-cultivated under the same set of growth conditions with only one varying environmental factor. Two competition pairs were investigated: (i) resistant mutants vs. the wild type and (ii) strongly resistant mutants (S1) vs. the other resistant mutants (S2, M1-3). Each revived strain was diluted in fresh growth media to reach the same OD_600_ (0.1). Then, two strains were mixed at a 1:1 (v/v) ratio and transferred at a 1:1000 dilution every 24 hours for three days (30 generations) under the designated growth conditions. The relatively low number of generations (30) was used to minimize the effect of adaptive mutations, a confounding factor that may alter the growth fitness. Co-cultures were sampled (∼ 700 μL) and centrifugated to collect cell pellets at the beginning and the end of the cultivation for the relative quantification by SNP genotyping assays. The remaining cultures were archived for further analysis. Triplicates were performed in each competition assay.

### 4.4. Determination of relative abundances of the two strains in a co-culture

Genomic DNA was extracted from cells collected from co-cultures using the DNeasy Blood and Tissue Kit (Qiagen) according to the manufacturer’s instructions. DNA concentration was determined by a Qubit 4 Fluorometer (Thermo Fisher Scientific, Wilmington, DE) and diluted to 3 ng/µL for subsequent qPCR-based SNP genotyping assay via Custom TaqMan SNP Genotyping Assays (Thermo Fisher Scientific). Those genotyping assays determined the relative abundance of each strain by calculating the ratios of a resistance gene allele to the wild type using specific primers and probes (**Table S2**). Assays were conducted in 96-well plates on a qPCR instrument QuantStudio 3, following the manufacturer’s recommended thermal cycling conditions (the details in the Supplementary Materials).

### 4.5. Determination of Relative Fitness

The relative fitness (F) of the resistant mutant was calculated using Equation 1 (Dykhuizen and Hartl, 1983; Lin et al., 2018), where R(t) and R_0_ represent the ratio of the resistant mutant to the competitor in the co-culture at time point t (R_t_) and the initial time (R_0_), respectively; n is the number of generations. With an initial ratio of 1:1 (R_0_ = 1), F > 1 means the ratio at time t increased and the resistant mutant had a growth advantage over its competitor, while F < 1 indicates a decreased ratio at time t and the growth of the resistant mutant was suppressed under the investigated growth condition.

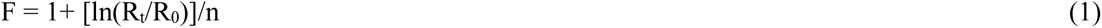

### 4.6. Statistical Analysis

Statistical analysis was performed using SPSS 19.0 (IBM, Armonk, USA). Differences of relative fitness between treatment and control groups were tested using Welch’s t-test with adjusted *p*-values using Bonferroni correction.

## CRediT authorship contribution statement

Y.X. and Y.M. designed the experiment; Y.X. and X.K. carried out the experiment; C.Z., Y.X., and X.K. analyzed the raw data; C.Z. and Y.X. summarized and visualized the results in figures; C.Z. and Y.M. drafted and revised the manuscript; all authors were involved in the results discussion and finalization of the manuscript.

## Acknowledgements

This study was supported by the National Science Foundation (Award No. 2045658).

## Supplementary Materials

Supplementary materials associated with this article can be found in the online version.

