## Supplementary Material for "Environmental Heterogeneity Altered the Growth Fitness of Antibiotic-Resistant Mutants and the Resistance Prevalence in *Escherichia coli* populations"

### Supplemental Methods

#### Preparation of M9 minimal media

The standard M9 minimal medium was prepared to a final volume of 1 L and included following components (Sambrook et al., 1989): 200 mL 5 × M9 salts stock solution (30 g/L $Na_2HPO_4$, 15 g/L $KH_2PO_4$, 5 g/L $NH_4Cl$, 2.5 g/L NaCl), 20 ml 20% (w/v) D-glucose (filter sterilized), 2 mL of 1 M $MgSO_4$, 0.1 mL of 1 M $CaCl_2$, and 1 mL trace element solution (16.67 g $FeCl_3 \cdot 6H_2O$, 0.18 g $ZnSO_4 \cdot 7H_2O$, 0.12 g $CuCl_2 \cdot 2H_2O$, 0.12 g $MnSO_4 \cdot 4H_2O$, 0.18 g $CoCl_2 \cdot 6H_2O$, and 22.5 g $Na_2EDTA \cdot 2H_2O$) (Fischer and Sauer, 2003). All components were sterilized separately: glucose and trace elements by filtration; salts and divalent cationic solutions by autoclavation. $CaCl_2$ was introduced after $MnSO_4$ to minimize precipitation.

**Table S1 M9 medium nutrient composition**

| Element | Source Compound(s) | Elemental conc. (mg/L) | Limited condition (fraction of the original) |
|---|---|---|---|
| C | D-glucose ($C_6H_{12}O_6$) | 1599.6 | 1/3 |
| N | $NH_4Cl$ | 262 | 2/3 |
| P | $Na_2HPO_4$, $KH_2PO_4$ | 1991.9 | 2/3 |
| S | $MgSO_4$, $MnSO_4 \cdot 4H_2O$, $ZnSO_4 \cdot 7H_2O$ | 64.2 | 1/2 |
| Ca | $CaCl_2$ | 4.0 | 1/100 |
| Mg | $MgSO_4$ | 48.6 | $1 \times 10^{-5}$ |
| Fe | $FeCl_3 \cdot 6H_2O$ | 3.4 | $1 \times 10^{-5}$ |

#### Genotyping assay

The genotyping reactions were carried out in 5-μL reaction mixtures containing 100 ng isolated RNA, 200 nmol/L each of the two primers, and 10 μL 2× SYBR green PCR mixture. The isolated mutants with the *rpsL, dsbC, glnE, nuoG548,* and *nuoG10* alleles confirmed by the SNP genotyping assay, as well as wild type, were used as controls. To quantify the fraction of strains in the pair, we set different fractions of the standard genomic DNA of the first strain to the standard genomic DNA of the second strain at 10%, 20%, 30%, 40%, 50%, 60%, 70%, 80%, and 90%. Three biological replicates were performed. The Thermo Fisher Cloud “Genotyping” application was used to generate allele calls. Primers used in this study are listed in **Table S2**.

**Table S2 Targeted mutant alleles and corresponding primers in qPCR-based SNP genotyping assay**

| | Forward primer sequence (5'→3') | Reverse Primer Sequence (5'→3') | VIC Probe Sequence (5'→3') | FAM Probe Sequence (5'→3') |
|---|---|---|---|---|
| *rpsL* | GAGCACTCCGTGAT CCTGATC | CGTACGGTGTGGT AACGAACA | CGGGAGGTCT TTAACAC | CGGGAGGTCTCT AACAC |
| *nuoG10* | GAAGCATGCTAATG GCTACAATTCA | GTTGTCCGCTCCGT TGAC | CGGCAAAGAA TACG | ACGGCAAATAAT ACG |
| *sbmA282* | GGAGTTTAAAAACC AGCGTGTAGAG | GCGTGGCATCGTCT TCAC | CATAAACCAG CTCTTTACG | AAACCAGCTATT TACG |
| *nuoG548* | CCCGTTCCGTCAAC AGCAT | CGGTTTCCAGTTCG GTTAACG | TCTTCAAGCG AACCGC | CTTCTTCAAGCTA ACCGC |
| *glnE* | TGACTGGCCGCAAC TGA | CTTTCATCGTCGCC AATCAATTCAT | TTGGTCATATG TGCGGTCAG | ATTGGTCATATGT ACGGTCAG |
| *dsbC* | AAAGCTATCTGGTG TGCGAAAGA | GCTGGTGCGACGC TTTT | CTGCCATCAC ATCAT | ACCTGCCATCTC ATCAT |

**Growth curve measurement**

*E. coli* strains were grown overnight to the stationary phase in 2 ml LB medium at 37 ºC. One µL of revived culture was transferred into 255 µL of LB media with designed experimental conditions on a 96-well plate, which was sealed with an opaque cover foil and incubated in a microplate reader (BioTek Instruments, Winooski, VT). Optical density was read at a wavelength of 600 nm ($OD_{600}$) automatically every 30 min for 24 h. The $OD_{600}$ of blank media was subtracted from the observed value before analysis. Eight replicates were performed. Due to edge effects on the 96-well plate, replicates with no obvious growth were treated as outliers and eliminated from the downstream analysis.

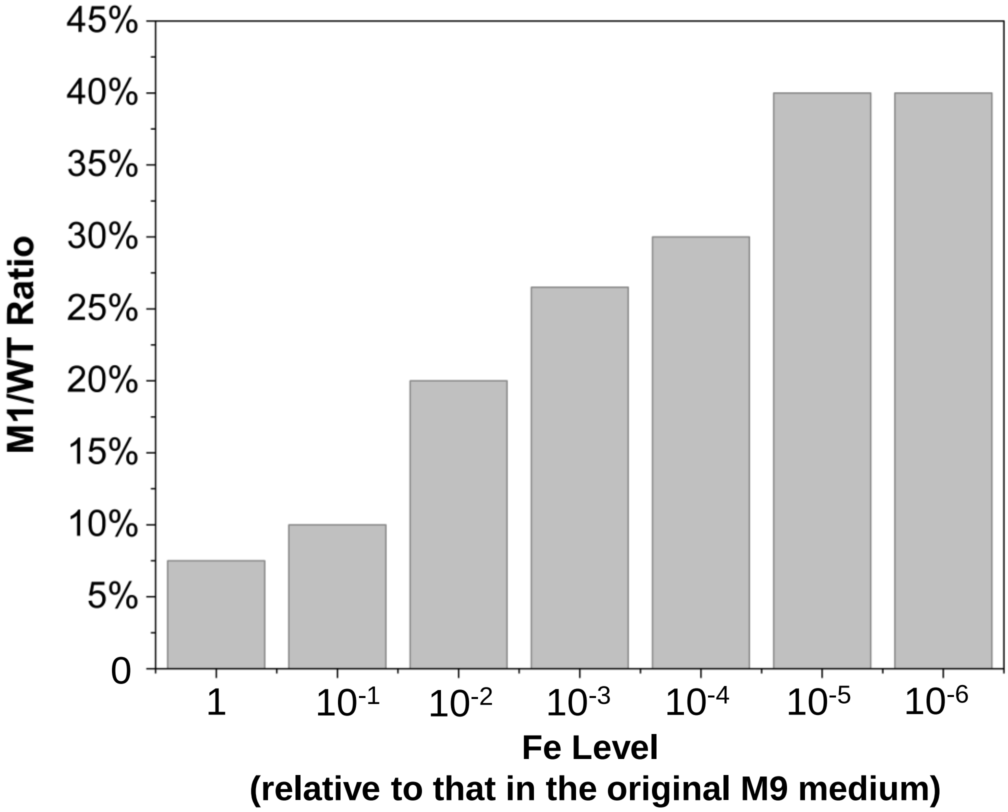


**Figure S1.** The relative abundance of the mildly resistant mutant M1 grown with the wild type in a co-culture in M9 medium with different Fe levels (1 – $10^{-6}$ of the concentration in the original M9 medium).

| With/without antibiotic stress | | Antibiotic Stress | | | | | | | |
|---|---|---|---|---|---|---|---|---|---|
| Competition pair | | RS1/RS2 | | RS1/RM1 | | RS1/RM2 | | RS1/RM3 | |
| Generation | | G0 | G30 | G0 | G30 | G0 | G30 | G0 | G30 |
| Temperature | 20 °C | 47% | 46% | 52% | 76% | 48% | 62% | 50% | 76% |
| | 30 °C | 47% | 51% | 52% | 93% | 48% | 74% | 50% | 94% |
| | 37 °C | 47% | 55% | 52% | 90% | 48% | 85% | 50% | 92% |
| | 45 °C | 47% | 53% | 52% | 95% | 48% | 94% | 50% | 97% |
| pH | pH = 5 | 47% | 30% * | 52% | 47% * | 48% | 25% * | 50% | 45% * |
| | pH = 6 | 47% | 41% | 52% | 53% * | 48% | 45% * | 50% | 55% * |
| | pH = 7 | 47% | 55% | 52% | 90% | 48% | 85% | 50% | 92% |
| | pH = 8 | 47% | 42% | 52% | 99% | 48% | 75% | 50% | 99% |
| Salinity_NaCl | 0 mM | 47% | 55% | 52% | 90% | 48% | 85% | 50% | 92% |
| | 50 mM | 47% | 50% | 52% | 90% | 48% | 85% | 50% | 85% |
| | 100 mM | 47% | 55% | 52% | 86% | 48% | 83% | 50% | 90% |
| | 200 mM | 47% | 47% | 52% | 90% | 48% | 90% | 50% | 95% |
| | 300 mM | 47% | 55% | 52% | 93% | 48% | 85% | 50% | 88% |
| Salinity_KCl | 0 mM | 47% | 55% | 52% | 90% | 48% | 85% | 50% | 92% |
| | 50 mM | 47% | 50% | 52% | 80% | 48% | 25% * | 50% | 45% * |
| | 100 mM | 47% | 50% | 52% | 65% | 48% | 15% * | 50% | 40% * |
| | 200 mM | 47% | 50% | 52% | 60% | 48% | 10% * | 50% | 20% * |
| | 300 mM | 47% | 50% | 52% | 50% * | 48% | 5% * | 50% | 5% * |

Fraction of RS1

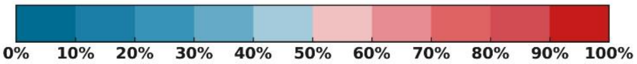


**Figure S2.** The ratio of S1 to other resistant mutants (S2, M1, M2, and M3) under various environmental conditions (i.e., temperature, pH, Na and K salinity) in the LB medium in the **presence** of selection pressure, i.e., 1/5 MIC streptomycin and a mixture of pesticides as described in the Materials and Methods section of the main text (the average values of triplicates are presented; asterisks indicate statistical difference from the positive control under optimal conditions, i.e., 37 °C, pH 7, and no salts added).
